# Identification of a Papain-Like Protease Inhibitor with Potential for Repurposing in Combination with an M^pro^ Protease Inhibitor for Treatment of SARS-CoV-2

**DOI:** 10.1101/2022.07.18.500363

**Authors:** Jesus Campagna, Barbara Jagodzinska, Pablo Alvarez, Constance Yeun, Patricia Spilman, Kathryn M. Enquist, Whitaker Cohn, Pavla Fajtová, Anthony J. O’Donoghue, Vaithilingaraja Arumugaswami, Melody M.H. Li, Robert Damoiseaux, Varghese John

## Abstract

SARS-CoV-2 requires two cysteine proteases for viral polypeptide processing to allow maturation and replication: the 3C-like protease also known as the Main protease (M^pro^) and the papain-like protease (PL^pro^). In addition to its critical role in viral replication, PL^pro^ removes post-translational modifications like ubiquitin and interferon-stimulated gene product 15 (ISG15) from host proteins through its deubiquitinase domain, leading to host immunosuppression and increased ability of the virus to evade the host antiviral immune response. Through screening of a custom clinical compound library, we identified eltrombopag (DDL-701), a thrombopoietin receptor agonist, as having PL^pro^ inhibitory activity that is sustained in the presence of the M^pro^ inhibitor nirmatrelvir. DDL-701 also suppressed both the deubiquitinase and ISG15 cleavage activities of PL^pro^. In addition, DDL-701 partially restored interferon-β induction – an element of the host immune response - in an *in vitro* model system. Further, modeling and docking studies suggest DDL-701 interacts with the active site region of the PL^pro^ enzyme and pilot pharmacokinetic studies indicate it is brain permeable. DDL-701 is already approved for treatment of thrombocytopenia and has previously been shown to achieve human plasma levels after oral dosing that is above the IC_50_ needed for it to exert its PL^pro^ inhibitory activity *in vivo*. In addition, it has also been reported to have antiviral efficacy against SARS-CoV-2. DDL-701 thus represents a drug that can immediately be repurposed and undergo clinical evaluation as a PL^pro^ inhibitor that may be most effectively used in a protease inhibitor cocktail with an M^pro^ inhibitor such as nirmatrelvir (Paxlovid) for the treatment of COVID-19.

## INTRODUCTION

Despite the development of vaccines, COVID-19 was the third-leading cause of death in the U.S. in 2021 [1] and new variants with the potential to evade vaccine protection have appeared, with more anticipated [2]. This will likely result in continued need for treatments that are effective post-infection. One strategy for development of antiviral medications to mitigate severe COVID-19 infections has been to target the two viral cysteine proteases, the 3C-like protease also known as the main protease (M^pro^) and the papain-like protease (PL^pro^), that are essential for polypeptide processing during viral maturation and replication [3-5].

The coronavirus nonstructural protein 3 (nsp3) has multifunctional domains including one of the two viral proteases used for initial cleavages of the coronavirus polyprotein through its PL^pro^ domain. In SARS-CoV, PL^pro^ has been reported to mediate cleavage of interferon (IFN) stimulated gene product 15 (ISG15) from interferon regulatory factor 3 (IRF3), blocking its nuclear translocation, and reducing type 1 IFN responses leading to host immune suppression [6, 7]. Nsp3 has several domains outside of the PL^pro^ domain including the nearby ubiquitin-like domain and macro domain [3]. It is thought that the ubiquitin-like domain works in concert with the PL^pro^ domain to carry out its de-ubiquitinating and de-ISGylating activities, which disrupt the host innate immune response by removing important ISG15 molecules that help promote the type I IFN response [8]. Additionally, the macro domain is thought to antagonize the innate immune response by removing ADP-ribose, a post-translational modification that enhances interferon signaling [6]. Previous studies have identified nsp3 as a main immune antagonist in many coronaviruses including HCoV-NL63, SARS-CoV, MERS-CoV, and SARS-CoV-2 [6, 7, 9-12]. Interestingly, both SARS and MERS viruses contain SARS-unique domains in nsp3 that enhance the innate immune antagonism activity of the PL^pro^ domain, which supports the merits of targeting the PL^pro^ for therapeutic treatment against SARS-CoV-2 [3, 13]. With the progression of COVID-19, studies have found a strong correlation between innate immune deficiency and severe outcomes [14] to which the host immune antagonism activity of nsp3 likely contributes.

In initial efforts to identify a therapeutic for SARS-CoV-2 infection, M^pro^ inhibitors were targeted, leading to the development and FDA emergency use authorization of nirmatrelvir (Paxlovid) [15]. M^pro^ is a promising drug target because it is dissimilar from human proteases and plays a critical role during infection. Most research has so far only focused on developing single inhibitors for M^pro^, and clinical data has shown that they successfully reduce hospitalization from COVID-19 [15].

Despite the benefits of nirmatrelvir treatment, there have been frequent reports of a ‘rebound’ effect [16, 17] that may be due, in part, to M^pro^ inhibition monotherapy not being completely effective in arresting SARS-CoV-2 replication. Recent studies suggest that the virus could develop resistance to nirmatrelvir through development of mutations in the protease M^pro^, a phenomenon observed with many antiviral drugs [18, 19]. This has raised interest in the development of protease inhibitor cocktail therapy for SARS-CoV-2 [20-27] to increase efficacy and reduce the risk of rebound, which is modeled on the use of protease cocktails for the treatment of HIV and hepatitis C. Use of a protease inhibitor cocktail could help reduce or prevent resistance by making it harder for the virus to evolve around multiple inhibitors.

Here, in pursuit of the objective of identifying a PL^pro^ inhibitor that may be used in a protease inhibitor cocktail with an M^pro^ inhibitor, we screened a custom 58-compound library of pharmacologically active molecules, including FDA-approved drugs, for PL^pro^ inhibition activity and identified the compound eltrombopag (DDL-701) that showed PL^pro^ inhibitory activity with an IC_50_ that is achieved in plasma after oral dosing. Importantly, the protease inhibition activities of both DDL-701 and nirmatrelvir are sustained when combined *in vitro*.

## RESULTS

### Screening of clinical compound library in PL^pro^ assay reveals hit DDL-701

We screened small molecules from a 58-compound custom clinical library that was assembled from commercial sources (see details in **Supplementary Table S1**) for their ability to inhibit PL^pro^ enzyme activity using a short peptide substrate. As shown in the scatterplot in **Figure 1A**, the screening revealed 4 hits that inhibit PL^pro^ activity > 80 %: eltrombopag (DDL-701), the known PL^pro^ inhibitor GRL-0617, thrombopoietin receptor agonist-1 (TPO-1 agonist, DDL-713), and zafirlukast (DDL-715) a potent cysteinyl leukotriene receptor antagonist [28]. Interestingly, two compounds – the antipsychotic drug fluspirilene and the leukotriene receptor antagonist montelukast - increased PL^pro^ activity (see compound well # in **Supplementary Table S1**).

**Figure 1.**
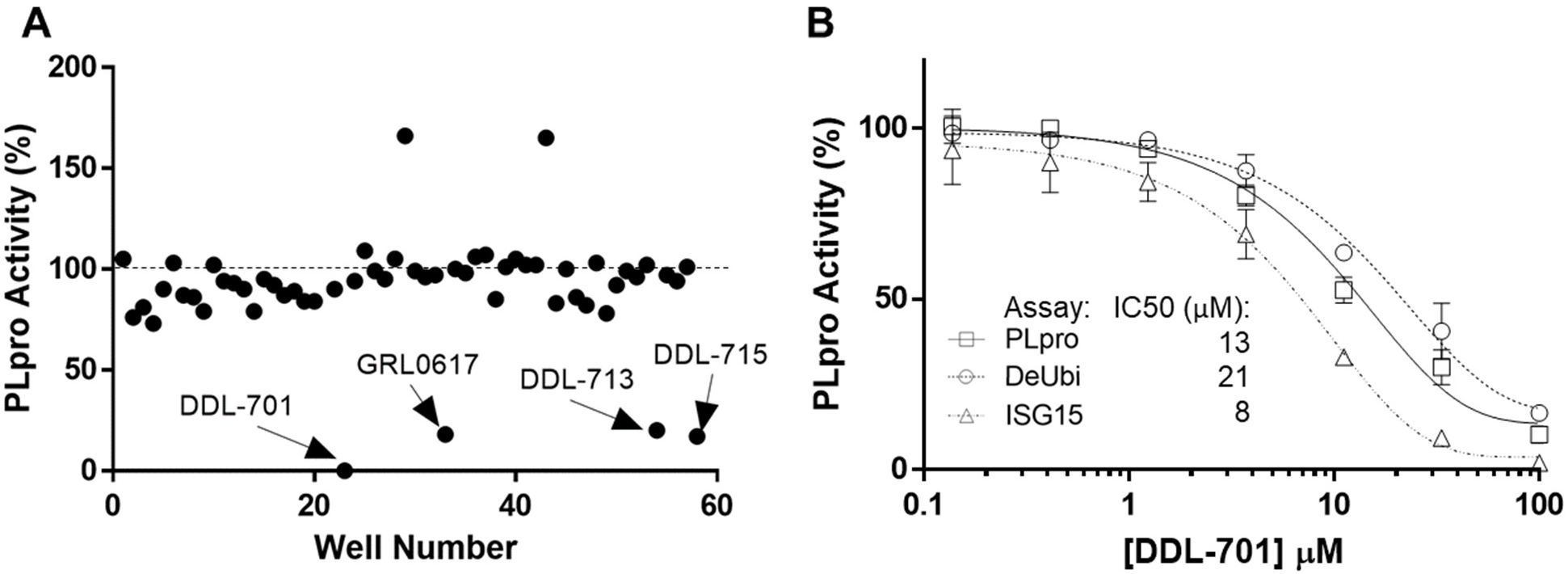
Identification of DDL-701 as a PL^pro^ inhibitor. (A) Scattergraph of clinical compound library screening using the PL^pro^ assay. (B) Dose-response and IC_50_ for DDL-701 in PL^pro^, deubiquitinase, and ISG15 assays.

To further validate DDL-701 as a PL^pro^ inhibitor, we assessed dose-response inhibition of PL^pro^, and its deubiquitinase and deISGylase activities (**Figure 1B**) which showed IC_50_ values of 13, 21, and 8 μM, respectively. Dose-response curves for PL^pro^ inhibition by the other hits from screening were also generated and displayed weaker potency than DDL-701: DDL-713 (IC_50PLpro_∼54 µM) and DDL-715 (IC_50PLpro_∼24 µM) (**Supplementary Figure S1**).

### DDL-701 and nirmatrelvir (DDL-750) show sustained protease activities in combination

For DDL-701 to be efficacious in protease cocktail therapy, it is important that its activity be sustained in the presence of an M^pro^ inhibitor such as nirmatrelvir (DDL-750). As shown in **Figure 2A**, inhibition of PL^pro^ by DDL-701 is not affected when it is used in combination with DDL-750 in the protease assay. Similarly, DDL-750 M^pro^ inhibitory activity was sustained in the presence of DDL-701 (**Figure 2B**). DDL-701 did not show activity in an M^pro^ activity assay.

**Figure 2.**
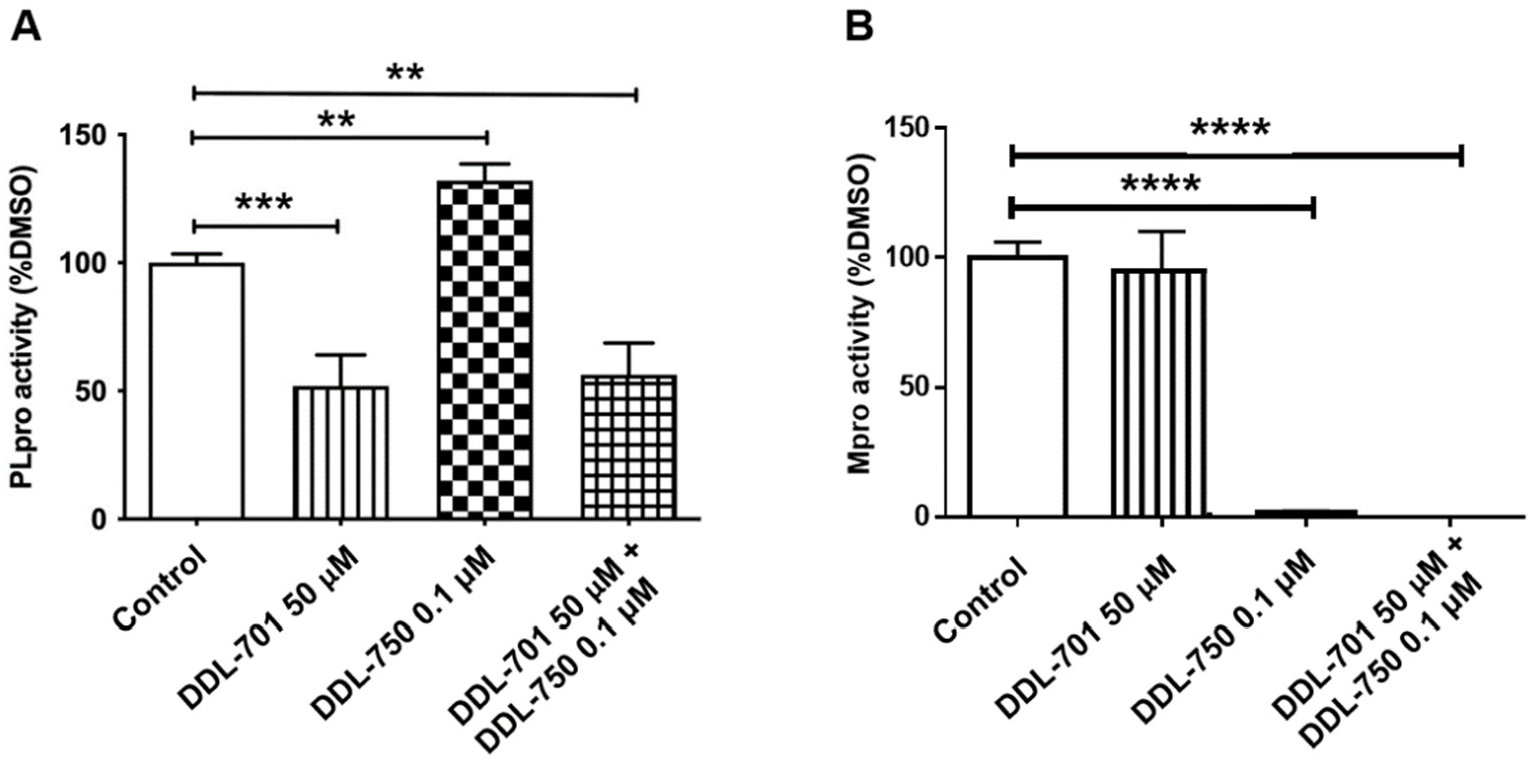
DDL-701 and DDL-750 alone and in combination in the PL^pro^ and M ^pro^ assays. The (A) PL^pro^ and (B) M^pro^ assays with DDL-701 and DDL-750 each alone and in combination at the concentrations shown. Statistical analysis performed using one-way ANOVA with post-hoc comparison of each treated well to the control, where ****p < 0.0001.

### DDL-701 activity partially restores interferon-β (IFN-β) induction in vitro

Because PL^pro^ activity is reported to inhibit the host cell IFN-β anti-viral response [6], we sought to determine if the PL^pro^ inhibitors can rescue IFN-β induction. HEK-293T cells were co-transfected with an IFN-β luciferase reporter and SARS-CoV-2 Wuhan strain PL^pro^, and stimulated with polyinosinic:polycytidylic acid (poly I:C) to mimic the double stranded RNA (dsRNA) that positive-sense, single-stranded RNA viruses such as SARS-CoV-2 form during viral replication. As shown in **Figure 3**, expression of PL^pro^ reduces IFN-β induction by > 50%, and treatment with DDL-701 at 1 µM modestly yet significantly rescued IFN-β induction (*p* = 0.042). Higher concentrations did not rescue induction. Neither DDL-715 nor DDL-750 rescued IFN-β induction (**Supplementary Figure S2A)**.

**Figure 3.**
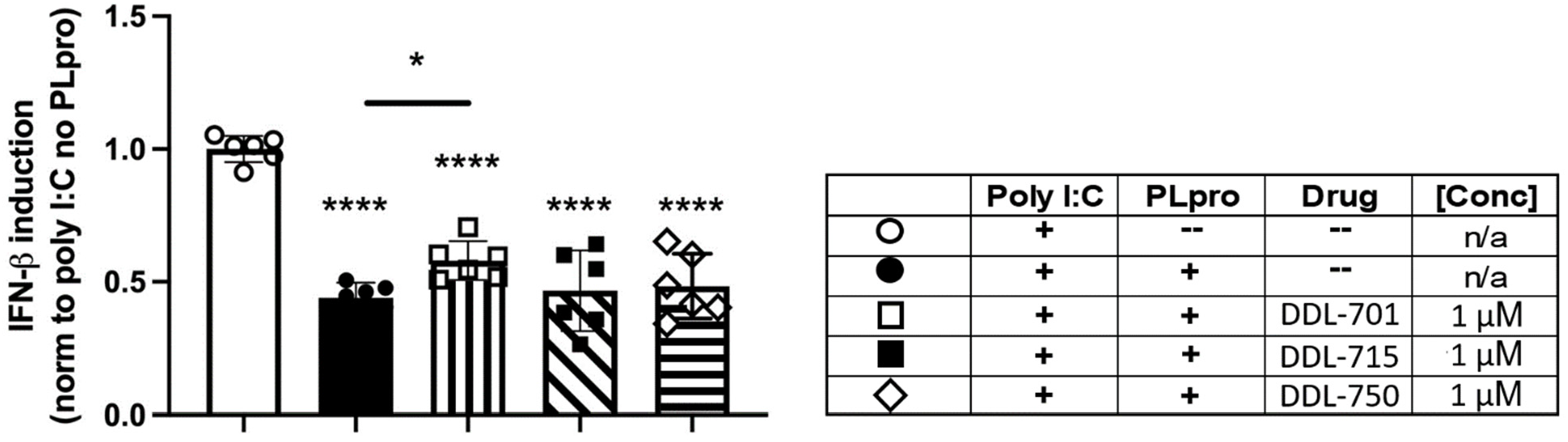
DDL-701 partially restores interferon-beta (IFN-β) induction by poly I:C in the presence of PL^pro^. Polyinosinic:polycytidylic acid (poly I:C) was used to stimulate IFN-β induction in HEK-293 cells. Transfection with a reporter construct expressing Wuhan strain PL^pro^ (see legend in table) significantly decreases IFN-β induction as compared to the poly I:C only control, which is partially restored by the PL^pro^ inhibitor DDL-701, but not by DDL-715 or M^pro^ inhibitor DDL-750, all at 1 μM. Statistical analysis performed using one-way ANOVA with Dunnett’s post-hoc comparison of the Poly I:C only control to all other groups and Sidak’s post-hoc comparison of Poly I:C + PL^pro^ no drug (--) to Poly I:C + PL^pro^ + DDL-701, where ****p < .0001 and *p ≤ .05.

PL^pro^ activity has also been reported to antagonize NFκB signaling [29], therefore an NFκB assay was also performed, but no rescue of NFκB signaling was observed by any compound tested under the conditions of the assay (**Supplementary Methods and Figure S2B**).

### DDL-701 is brain-permeable in mice

To determine if DDL-701 as part of a possible protease inhibitor cocktail has the potential to be effective against SARS-CoV-2 infection of the central nervous system (CNS), a pharmacokinetic study was performed in mice. Animals received DDL-701 or -750 alone or in combination both at 30 mg/Kg by oral dosing, and brain and plasma levels assessed. As shown in **Figure 4**, 2 hours after oral treatment, brain levels of DDL-701 were ∼1 μM (∼442 ng/g) while DDL-750 (nirmatrelvir) reached a concentration of ∼0.18 μM (∼91 ng/g). DDL-701 has >5 fold higher brain penetration than nirmatrelvir. In plasma, DDL-701 and DDL-750 levels were ∼14 μM (∼ 6,214 ng/mL) and ∼0.5 μM (∼234 ng/mL) respectively. In combination, the levels in plasma and brain were lower for both drugs; DDL-701 levels in plasma were ∼4 μM (∼1,628 ng/mL) and in brain ∼0.4 μM (∼ 169 ng/g), and DDL-750 levels were ∼0.1 μM (∼55 ng/mL) and ∼0.1 μM (∼ 59 ng/g) in plasma and brain, respectively.

**Figure 4.**
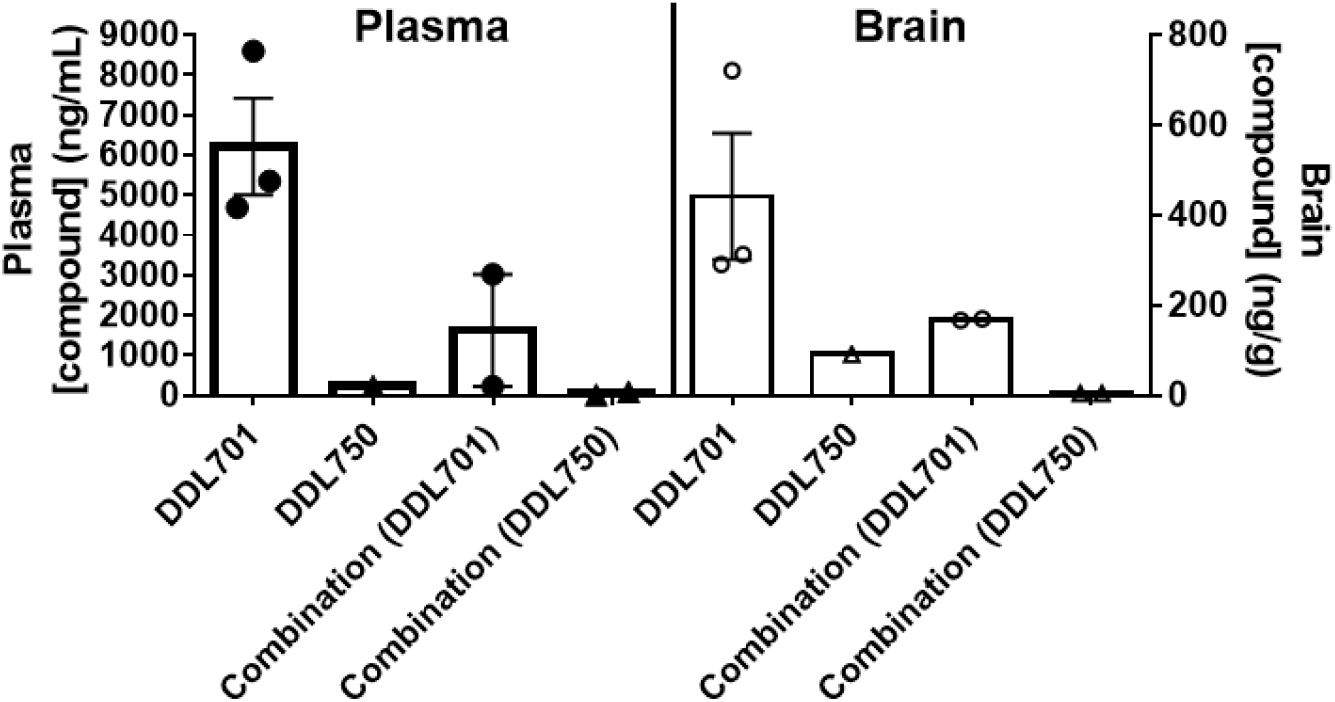
Plasma and brain levels of DDL-701 and DDL-750 in mice. Mice received either DDL-701 or 750 alone or in combination by oral dosing at 30 mg/Kg. Plasma (left) and brain (right) levels were assessed 2 hours after dosing. DDL-701 n = 3, DDL-750 n = 1, and DDL-701 + 750 n = 2.

### Modeling and docking reveal DDL-701 and known PL^pro^ inhibitor GRL-0617 interact with residues near the active site

Modeling by docking of DDL-701 with PL^pro^ was compared with that of the known PL^pro^ inhibitor GRL-0617 (PDB ID: 7CMD) [30] using Cresset software. The docking data suggests that neither GRL-0617 or DDL-701 (**Figure 5A and B**, respectively) interacts with the active site cysteine-111 residue, but rather are bound to the site around tyrosine-268 which lies outside the tunnel containing the active site residue Cys111. DDL-715 and the reported PL^pro^ inhibitor losartan [22] bind the enzyme similarly to DDL-701, at the site around Tyr268 and at the entrance of the active site tunnel leading to Cys111 (**Supplementary Figures S3 and S5, respectively**). Similarly, docking of DDL-713 which is a TPO agonist like DDL-701 (**Supplementary Figure S4**) binds around Tyr268 located outside the catalytic-site cavity containing the key residue Cys111.

**Figure 5.**
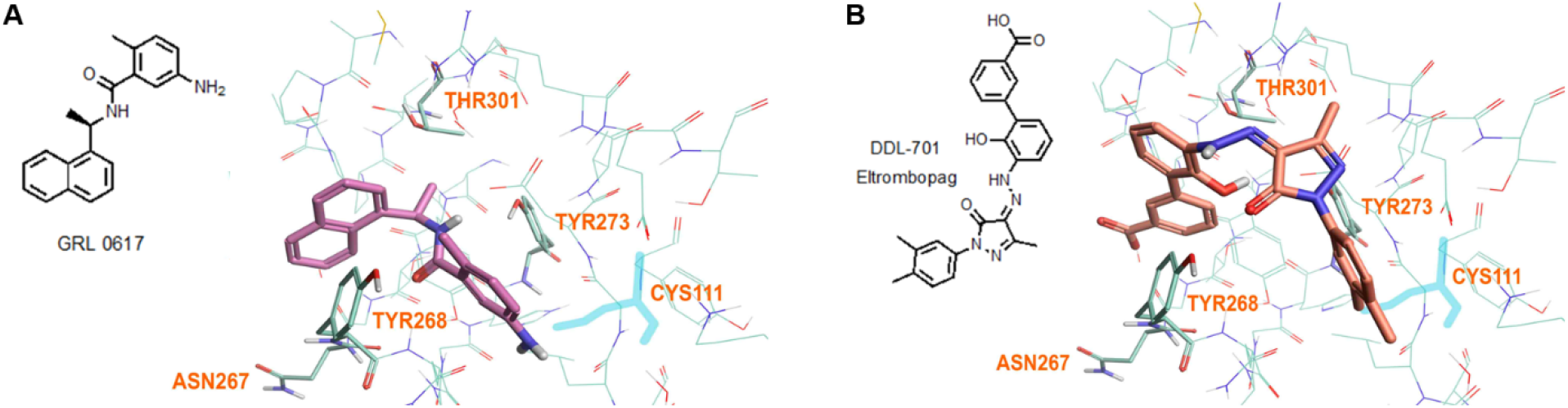
Model of GRL-0617 (PL^pro^ inhibitor) and DDL-701 (Eltrombopag) in the active site of PL^pro^. (A) X-ray crystallography structure of PL^pro^ and GRL-0617 (PDB ID: 7CMD) is shown. (B) DDL-701 binds similar to GRL-0617, based on the PL^pro^ amino acid residues marked in the vicinity these two ligand binding sites. Both molecules bind at the entrance of the active site tunnel around Tyr268 and distal from the active site residue Cys111.

## DISCUSSION

DDL-701, identified in our screening of a library of clinical compounds, shows potent PL^pro^ inhibition and elicits partial restoration of IFN-β induction *in vitro*. DDL-701 has an advantage for development because it is already approved for thrombocytopenia (available in the US under the brand name PROMACTA and outside the US as REVOLADE), and human pharmacokinetic (PK) data is available. These reported PK data indicate that a single oral dose of 75 mg DDL-701 in tablet form results in plasma C_max_ levels in the range of 25-30 μM with an AUC of 168 μg-hour/mL [31], a level that is above the IC_50_ of ∼ 13 μM for inhibition of PL^pro^. DDL-701 thus would be likely to exert PL^pro^ inhibition *in vivo* in human patients after an oral dose.

Another PL^pro^ inhibitor identified from our screening, DDL-715, is less potent than DDL-701 and in human achieves a C_max_ of ∼ 0.5 μM in plasma after oral dosing [28]. This is similar to what was shown with another PL^pro^ inhibitor losartan that also has a reported IC_50_ of ∼13 μM for PL^pro^ inhibition [21]. After oral dosing with losartan, however, the C_max_ in plasma is only ∼ 0.5 μM, well below its IC_50_ for PL^pro^ inhibition. Losartan, in our modeling, also bound the active site region similarly to DDL-701 (**Supplementary Fig S5**). The PL^pro^ inhibitor tropifexor, a potent Farnesoid X Receptor agonist, is reported to have a PL^pro^ IC_50_ ∼6 μM [25]. The binding of tropifexor in our modeling is similar to DDL-701 (**Supplementary Figure S6**). The C_max_ in human plasma with an approved oral dose is <0.1 μM, well below its PL^pro^ IC_50_. All of these data show that DDL-701 has an advantage, in that after a clinically used oral dose of the drug it is possible to achieve plasma levels above the PL^pro^ IC_50,_ making it a promising candidate to repurpose as part of the protease inhibitor cocktail with a M^pro^ inhibitor such as nirmatrelvir for treatment of COVID-19.

Here, we also report that our pilot PK studies in mice show DDL-701 is brain-permeable, which suggests it has potential to counter the effects of SARS-CoV-2 infection and potential complications in the CNS [32]. Nirmatrelvir had lower brain-permeability, but it is highly potent for the M^pro^ enzyme with an IC_50_ of ∼3 nM [15], therefore it may be effective in brain as part of the protease inhibitor cocktail. The establishment of such efficacy awaits testing in animal models of SARS-CoV-2 infection and/or in clinical trials.

As further evidence of its potential, DDL-701 has also previously been reported to have antiviral efficacy in Vero E6 cells infected with SARS-CoV-2 with an EC_50_ of 8 μM [20], although a mechanism of action for eltrombopag in that report is not described. In a separate study, eltrombopag was reported to show the ability to bind and reduce the stability of the spike protein-ACE2 complex [33], which may contribute to its antiviral efficacy. Importantly, the plasma C_max_ level of DDL-701 after a single oral dose in humans is well above this EC_50_ for inhibition of SARS-CoV-2 infection.

Ideally, the PL^pro^ inhibitor DDL-701 used in a protease inhibitor cocktail with the M^pro^ nirmatrelvir would result in induction of host antiviral activity, in addition to reducing viral polypeptide processing and replication. During viral infection there is induction of host antiviral response, cytosolic viral RNA forms a complex with the proteins RIG-1/MDA5, interacts with the molecular cascade involving MAVS, TBK1 and IRF3, leading to phosphorylation and nuclear translocation of IRF3 inducing transcription of type I interferons [6], this response can be antagonized by PL^pro^ (**Figure 6**). By interfering with PL^pro^ cleavage of ISG15 and thus IRF3 phosphorylation [34], DDL-701 may restore IFN-β signaling *in vivo*, a potential effect supported by our data showing increased IFN-β induction *in vitro*. Confirmation of this potential mechanism awaits further study.

**Figure 6.**
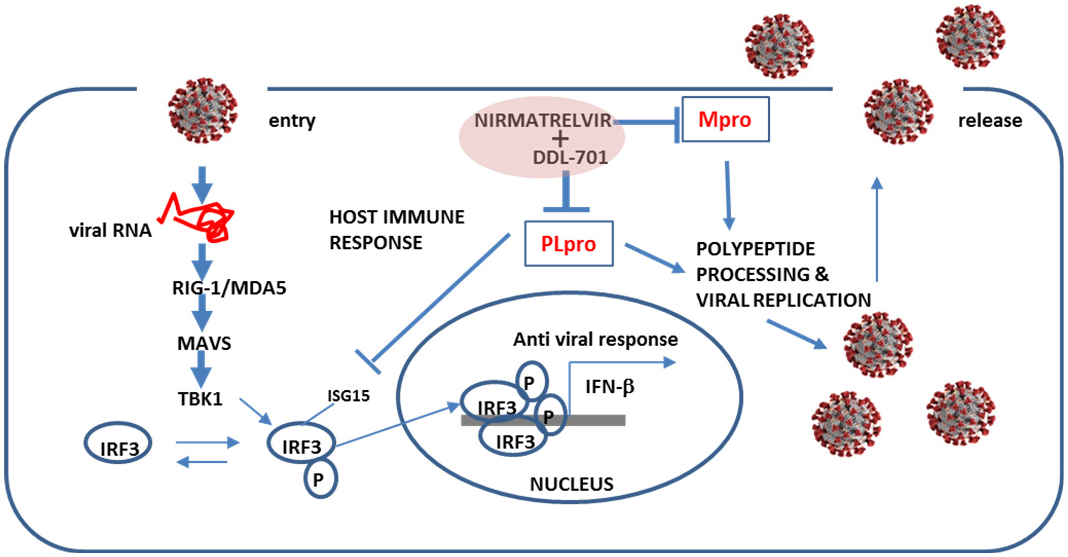
Putative restoration of host immune responses by a DDL-701/nirmatrelvir cocktail. PL^pro^ alters IRF3 phosphorylation, activity and the IFN-β host response. In combination with an M^pro^ inhibitor such as nirmatrelvir, it could help restore the host cell immune response in addition to reducing viral replication.

While the SARS-CoV-2 virus does not easily enter the brain, it has been reported to damage endothelial cells in the blood brain barrier (BBB) leading to inflammation and brain injury [35]. Knowledge concerning the impact of SARS-CoV-2 infection on the CNS and cerebrovasculature is limited and still being elucidated. The entry of the virus into the brain through the olfactory nerve endings and at the blood-CSF barrier (choroid plexus) is postulated [32, 36]. Evidence also suggests that the virus can infect astrocytes and could lead to some of the observed neurological symptoms such as fatigue, depression and brain fog [35]. Our pilot pharmacokinetic studies in mice herein suggest DDL-701 is brain-permeable, and while the brain permeability of nirmatrelvir is low, given its nanomolar potency in inhibiting viral replication [15] as part of the protease inhibitor cocktail it may have the potential to lower the risk of CNS complications.

Based on our findings that DDL-701 is a potent PL^pro^ inhibitor with sustained inhibitory activity in the presence of M^pro^ inhibitor nirmatrelvir and has previously been reported to achieve plasma levels that are likely to elicit both PL^pro^ inhibition and antiviral efficacy. It is a promising candidate for study in combination with a M^pro^ inhibitor as a protease inhibitor cocktail for the treatment of SARS-CoV-2 infection. DDL-701’s brain permeability suggested by our studies further points its potential to reduce infection in the CNS. DDL-701 has the advantage of being a re-purposed drug already available in an oral tablet that could be re-formulated with oral Paxlovid. Such a protease inhibitor cocktail presents the possibility of a more effective treatment for patients with COVID-19 that has the potential to reduce hospitalization and deaths resulting from the disease, speed up post-infection recovery, and reduce the risk of rebound, antiviral resistance, and CNS complications.

## METHODS

### PL^pro^, M^pro^, deubiquitinase, and ISG15 in vitro assays

PL^pro^ and deubiquitinase kits were purchased from BPS Bioscience (cat# 79995-2, cat# 79996, respectively), and for the ISG15 assay, rhodamine 110 was purchased from South Bay Bio (cat# SBB-PS0002); each assay was performed following the manufacturer’s recommendations, adapted to a 384-well plate format. For the M^pro^ assay, the fluorogenic substrate for SARS-CoV-2 M^pro^ was purchased from Vivitide (cat# SFP-3250-v). Briefly, in each assay, the enzyme was loaded into each well in the appropriate buffer, next the compounds were added into the well and incubated for 10-60 min at 37°C. The reaction was initiated by addition of substrate to each well and the fluorescent signal was read for 60 min at the appropriate excitation/emission. For the scatterplot analysis of the custom clinical library compounds treatment was done at 50 μM. For the M^pro^ activity assay, 8 µL of M^pro^ at 5 µg/mL in assay buffer was loaded into each well, followed by 100 nL of the compounds at 50 μM and incubated for 10 min at 37°C. Following the incubation, 2 µL of substrate at 25 µM was added into the wells and the fluorescent signal (ex/em 380/455nm) was recorded for 1 h at 37°C.

### Cloning of expression and reporter constructs

The SARS CoV-2 papain-like protease (PL^pro^) domain of Nsp3 was cloned from a doxycycline-inducible piggyBac transposon vector (PB-TAC-ERP2, Addgene# 80478) containing the synthesized full-length Nsp3 from the Wuhan-Hu-1 SARS CoV 2 strain (Alvarez and Yao, unpublished). The PL^pro^ domain (amino acids 745-1061) was cloned using PCR (Forward primer 5’-GTTTGCGGCCGCAAGAGAGGTGAGAACCATCAAG-3’, Reverse primer 5’-GGTGTCTAGATTAAGGTTTTATGGTTGTGGTATAG-3’) and inserted into myc-pcDNA3.1 using NotI and XbaI restriction sites. For the *Renilla* luciferase reporter construct, the *Renilla* luciferase gene was amplified from a Japanese encephalitis virus replicon containing a *Renilla* luciferase reporter gene using PCR (Forward primer 5’-GTTTAAGCTTGCCACCATGGCTTCCAAGGTGTAC-3’, Reverse primer 5’-TTTGCTCGAGTCACTGCTCGTTCTTCAGCAC-3’) and inserted into V5-pcDNA3.1 using HindIII and XhoI restriction sites, removing the V5 tag. An Addgene plasmid was used for the IFN-β promoter luciferase reporter (IFN-Beta_pGL3, #102597). To make the NF-κB signaling reporter, a gene block containing 5 tandem consensus NF-κB binding sites (Badr et al 2009; Ngo et al 2020), followed by the -55 to +19 region of the human IFN-β gene (UCSC Genome Browser) was synthesized by Integrated DNA Technologies (Alvarez and Yao, unpublished). The gene block was then moved into the pCR™-Blunt II-TOPO™ vector using the Zero Blunt™ TOPO™ PCR Cloning Kit (ThermoFisher) and using EcoRI and NheI to move the gene block into IFN-Beta_pGL3, replacing the IFN-β promoter.

For SARS CoV-2 M^pro^, the plasmid was provided by Rolf Hilgenfeld, University of Lübeck, Germany [37] and transformed into Escherichia coli strain BL21-Gold (DE3). The expression and purification of the protein has been described in detail previously [38].

### IFN-β induction and NFκB signaling luciferase reporter assays

To assess whether the compounds could rescue IFN-β induction, HEK293T cells were treated with compound by replacing the media with new media containing the compound at the desired concentration. The cells were then co-transfected with 150 ng of an IFN-β promoter-*Firefly* luciferase reporter plasmid, 10 ng of a *Renilla* luciferase expression plasmid, and 250 ng of either an empty myc-pcDNA3.1 vector or a plasmid expressing SARS-CoV-2 PL^pro^ domain of Nsp3 (amino acids 745-1061), using the TransIT-mRNA Transfection Kit (Mirus bio). To induce IFN-β, 500 ng of poly I:C (InvivoGen) was added into the co-transfection mix.

For the NFκB signaling assay, cells were co-transfected with 150 ng of an NFκB dependent-*Firefly* luciferase reporter plasmid wherein the luciferase gene expression was under control of 5 tandem NFκB binding sites followed by the -55 to +19 region of the human IFNB1 gene for correct spacing, 10 ng of a *Renilla* luciferase expression plasmid, and 250 ng of either an empty myc-pcDNA3.1 vector or a plasmid expressing SARS-CoV-2 PL^pro^ domain of Nsp3 (amino acids 745-1061), using the TransIT-mRNA Transfection Kit (Mirus bio). To stimulate signaling, cell media was replaced with media containing the compound and 20 ng/mL tumor necrosis factor alpha TNF-α (R&D Systems).

For both assays, cells were harvested 24 hours post transfection using 1X passive lysis buffer and the dual luciferase assay was performed according to instructions from manufacturer (Promega). Luciferase signals were read using a BioTek Synergy H1 plate reader (Thermo Fisher Scientific).

### Pharmacokinetics analysis in mice

Adult C57BL/6 mice received DDL-701 or DDL-750 in DMSO alone or in combination at 30 mg/Kg (each) by oral gavage. Two hours after dosing, mice were anesthetized and blood was collected following transcardial puncture for the isolation of plasma. Mice were then perfused transcardially with saline and brain tissue removed. Levels of DDL-701 and DDL-750 were determined in brain and plasma by liquid chromatography-tandem mass spectrometry (LC-MS/MS). Briefly, a targeted LC-MS/MS assay was developed for each compound using the multiple reaction monitoring (MRM) acquisition method on a 6460 triple quadrupole mass spectrometer (Agilent Technologies) coupled to a 1290 Infinity HPLC system (Agilent Technologies) with a Phenomenex analytical column (Kinetex 1.7 µm C18 100 Å 100 × 2.1 mm). The HPLC method utilized a mixture of solvent A (99.9/1 Water/Formic Acid) and solvent B (99.9/1 Acetonitrile/Formic Acid) and a gradient was use for the elution of the compounds (min/%B: 0/20, 3/20, 19/99, 20/99, 21/20, 35/20). In this assay, detection of fragmented ions originating from each compound at specific chromatographic retention times were utilized to ensure specificity and accurate quantification in the complex biological samples (DDL-701 (M+H)^+^: 443.1, fragment ions: 211.0/322.1, retention time: 21.6; DDL-750 (M+H)^+^: 500.2, fragment ions: 111.0/319.1, retention time: 9.1). Chromatographic peak areas from standards made in drug naïve plasma and brain tissue lysates with increasing amounts of DDL-701 and DDL-750 were used to make a standard curve, and the trendline equation was used to calculate the absolute concentrations of each compound.

### Modeling of compound docking to the PL^pro^ active site

The crystal structure of PL^pro^ protein (PDB ID: 7CMD) with GRL-0617 ligand [39] was used as a basis for docking studies. The binding site was defined by placing GRL-0617 in the center of the grid box of approximately 20Å dimensions.

Flare (Cresset Software, v5.0.0) was used for docking modeling. The protein was prepared for docking utilizing an internal optimization module. Three separate docking runs were performed for each compound, each generating ∼200 poses, and each set of poses was evaluated for the best fit. For the final manual expert evaluation, ∼5-10 poses were selected and then further narrowed to 1-2 poses with the best scores. Each final pose was then examined to identify likely interactions with protein residues (hydrogen bonding, π-π or hydrophobic interactions, etc.). Orientations of compounds showed strong structural interaction with Tyr268 that lies outside the active site tunnel and were considered as preferred binding site of these molecules through π-π and hydrogen bonding interaction with Tyr268. Such compound binding interaction could interfere with PL^pro^ substrate binding and proteolytic activity.

## Supporting information

Campagna PLpro Inhib Supplementary Materials

## ACKNOWLEDGEMENTS

We would like to thank the David Geffen School of Medicine at UCLA for the pilot grant (OCRC #20-69) to VJ and the pilot grant (OCRC #20-16) to RD, the CNSI and the DDL that enabled the screening and identification of the drug candidates. ML was supported in part by the Johanna and Joseph H. Shaper Family Chair.

## REFERENCES

1. Soneji S. Population-level mortality burden from novel coronavirus (COVID-19) in Europe and North America. Genus 2021; 77(1):7. DOI:10.1186/s41118-021-00115-9.

2. Li X. Omicron: Call for updated vaccines. J. Med. Virol. 2022; 94(4): 1261–1263. DOI: 10.1002/jmv.27530.

3. Lei J, Kusov Y & Hilgenfeld R. Nsp3 of coronaviruses: Structures and functions of a large multi-domain protein. Antiviral Res. 2018; 149, 58–74. DOI: 10.1016/j.antiviral.2017.11.001

4. Ratia K et al. Structural Basis for the Ubiquitin-Linkage Specificity and deISGylating Activity of SARS-CoV Papain-Like Protease. PLoS Pathog. 2014; 10, e1004113. DOI: 10.1371/journal.ppat.1004113

5. Yadav R et al., Role of Structural and Non-Structural proteins and therapeutic targets and SARS-CoV-2 for COVID-19. Cell 2021; 10(4):821. DOI: 10.3390/cells10040821.

6. Shin D. et al. Papain-like protease regulates SARS-CoV-2 viral spread and innate immunity. Nature 2020; 587, 657–662. DOI: 10.1038/s41586-020-2601-5.

7. Matthews K., et al. The SARS coronavirus papain-like protease can inhibit IRF3 at a post activation step that requires deubiquitination activity. Virol. J. 2014; 11, 209. DOI: 10.1186/s12985-014-0209-9.

8. Freitas B, et al. Characterization and Noncovalent Inhibition of the Deubiquitinase and deISGylase Activity of SARS-CoV-2 Papain-Like Protease. ACS Infect. Dis. 2020; 6, 8, 2099–2109. DOI: 10.1021/acsinfecdis.0c00168.

9. Li C, et al. Viral Macro Domains Reverse Protein ADP-Ribosylation. J. Virol. 2016; 90, 8478–8486. DOI: 10.1128/JVI.00705-16.

10. Malgras M, et al. The Antiviral Activities of Poly-ADP-Ribose Polymerases. Viruses 2021; 13, 582. DOI: 10.3390/v13040582.

11. Sun L, et al. Coronavirus Papain-like Proteases Negatively Regulate Antiviral Innate Immune Response through Disruption of STING-Mediated Signaling. PLoS ONE 2012; 7, e30802. DOI: 10.1371/journal.pone.0030802.

12. Mielech A M, et al. MERS-CoV papain-like protease has deISGylating and deubiquitinating activities. Virology 2014; 450–451, 64–70. DOI: 10.1016/j.virol.2013.11.040.

13. Ma-Lauer Y, et al. p53 down-regulates SARS coronavirus replication and is targeted by the SARS-unique domain and PL pro via E3 ubiquitin ligase RCHY1. (2016) Proc Natl Acad Sci U S A. 2016 Aug 30;113(35):E5192–201. DOI: 10.1073/pnas.1603435113.

14. Hadjadj J, et al. Impaired type I interferon activity and inflammatory responses in severe COVID-19 patients. Science 2020; 369, 718–724. DOI: 10.1126/science.abc6027.

15. Owen DR, et al. An oral SARS-CoV-2 Mpro inhibitor clinical candidate for the treatment of COVID-19. Science 2021; 374(6575):1586–1593. DOI: 10.1126/science.abl4784.

16. Gupta K, et al. Rapid relapse of symptomatic omicron SARS CoV-2 infection following early suppression with nirmatrelvir/ritonavir. 26 April 2022, PREPRINT (Version 1) available at Research Square [https://doi.org/10.21203/rs.3.rs-1588371/v1].

17. Carlin AF, et al. Virologic and Immunologic Characterization of COVID-19 Recrudescence after Nirmatrelvir/Ritonavir Treatment. 18 May 2022, PREPRINT (Version 1) available at Research Square [https://doi.org/10.21203/rs.3.rs-1662783/v1].

18. Sacco MD, et al. The P132H mutation in main protease of omicron SARS-CoV-2 decreases thermal stability without compromising catalysis or small-molecule drug inhibition. Cell Res 2021; 32, 498–500. DOI: 0.1038/s41422-022-00640-y.

19. Zhou Y, et al., Nirmatrelvir Resistant SARS-CoV-2 Variants with High Fitness In vitro. BioRxiv 2022; DOI:.10.1101/2022.06.06.494921.

20. Jeon S, et al. Identification of Antiviral Drug Candidates against SARS-CoV-2 from FDA-Approved Drugs. Antimicrob Agents Chemother. 2020; 64(7): e00819–20. DOI: 10.1128/AAC.00819-20.

21. Nejat R et al., Losartan inhibits SARS-CoV-2 Replication in vitro. J. Pharm Pharm Sci. 2021; 24, 390–399. DOI: 10.18433/jpps31931.

22. Jamir E, et al. Applying polypharmacology approach for drug repurposing for SARS-CoV-2. J. Chem. Sci. 2022, 134;57. DOI: 10.1007/s12039-022-02046-0.

23. Lewis DSM et al., Aloin isoforms (A and B) selectively inhibits proteolytic and deubiquinating activity of PLpro of SARS-CoV-2 in vitro. Sci. Rep. 2022; 12, 2145. DOI: 10.1038/s41598-022-06104-y.

24. Napolitano V, et al., Acriflavine, a clinically approved drug, inhibits SARS-CoV-2 and other betacoronaviruses. Cell Chem Bio 2011; 29, 774–784. DOI: 10.1016/j.chembiol.2021.11.006.

25. Chunlong M, et al., Drug-Repurposing Screening Identified Tropifexor as a SARS-CoV-2 Papain-like Protease Inhibitor. ACS Infectious Disease 2022; 8, 1022. DOI: 10.1021/acsinfecdis.1c00629.

26. Narayanan A, et. al., Identification of SARS-CoV-2 inhibitors targeting Mpro and PLpro using in cell protease assay. Commun. Biol. 2022; 5, 169. DOI: 10.1038/s42003-022-03090-9.

27. Kulandaisamy R, et al., Repurposing of FDA approved Drugs against SARS-CoV-2 papain-like protease: Computational, Biochemical and in vitro studies. Front. Microbiol. 2022; 13:877813. DOI: 10.3389/fmicb.2022.877813.

28. Murata Y & Sugimoto O. Zafirlukast (Accolate): a review of its pharmacological and clinical profile. Nihon Yakurigaku Zasshi. 2002 Apr;119(4):247–58. DOI: 10.1254/fpj.119.247.

29. Frieman M, et al., Severe acute respiratory syndrome coronavirus papain-like protease ubiquitin-like domain and catalytic domain regulate antagonism of IRF3 and NF-kappaB signaling. J. Virol. 2009;83(13):6689–705. DOI: 10.1128/JVI.02220-08.

30. Gao X, et al., Crystal structure of SARS-CoV-2 papain-like protease. Acta Pharm Sin B. 2021; 11: 237–245. DOI: 10.1016/j.apsb.2020.08.014.

31. Matthys et al., Clinical pharmacokinetics, platelet response, and safety of eltrombopag at supratherapeutic doses of up to 200 mg once daily in healthy volunteers. J Clin Pharmacol. 2011 Mar;51(3):301–8. DOI: 10.1177/0091270010368677.

32. Gupta, M and Weaver D. COVID-19 as a trigger of Brain Autoimmunity. ACS Chem Neurosci. 2021; 12:2558. DOI: 10.1021/acschemneuro.1c00403.

33. Feng S et al. Eltrombopag is a potential target for drug intervention in SARA-CoV-2 spike protein. Infect Genet Evol. 2020, 85: 104419. DOI: 10.1016/j.meegid.2020.104419.

34. GuanQun L, et al., ISG15-dependent activation of the sensor MDA5 is antagonized by the SARS-CoV-2 papain-like protease to evade host innate immunity. Nat Microbiol. 2021, 6: 467–468. DOI: 10.1038/s41564-021-00884-1.

35. Boldrini M, et al. How COVID-19 affects the brain. JAMA Psychiatry 2021;78(6):682–683. DOI:10.1001/jamapsychiatry.2021.0500.

36. McQuaid C, et al., SARS-Cov-2: is there neuroinvasion? Fluids Barriers CNS 2021; 18: 32. DOI: 10.1186/s12987-021-00267-y.

37. Zhang L, et al., Crystal structure of SARS-CoV-2 main protease provides a basis for design of improved ?-ketoamide inhibitors. Science 2020; 368(6489): 409–412. DOI: 10.1126/science.abb3405.

38. Mellott DM, et al., A clinical-stage cysteine protease inhibitor blocks SARS-CoV-2 infection of Human and Monkey Cells. ACS Chem. Biol. 2021, 16, 642–650. DOI: 10.1021/acschembio.0c00875.

39. Fu Z et al. The complex structure of GRL0617 and SARS-CoV-2 PLpro reveals a hot spot for antiviral drug discovery. Nat Commun. 2021;12: 488. DOI: 10.1038/s41467-020-20718-8.

