## Supplementary material for "Identification of a Papain-Like Protease Inhibitor with Potential for Repurposing in Combination with an M^pro^ Protease Inhibitor for Treatment of SARS-CoV-2": Campagna PLpro Inhib Supplementary Materials

### Supplementary Figures

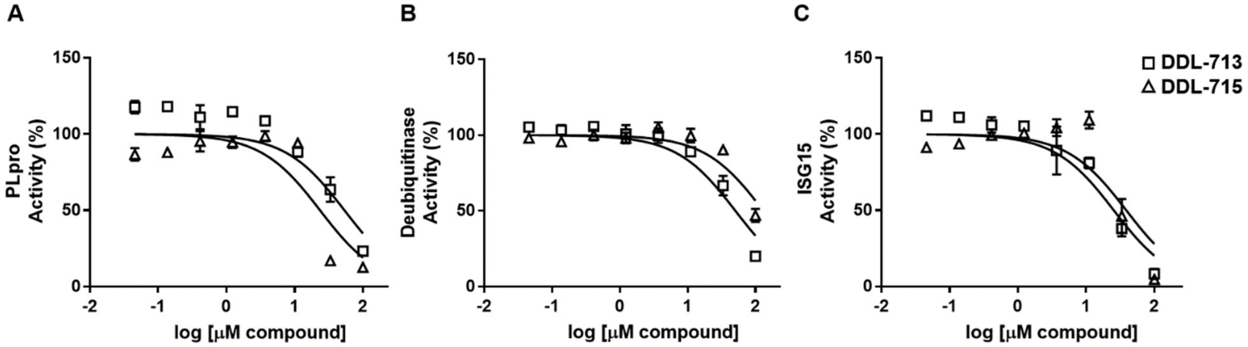


**Figure S1**. *DDL-713 and DDL-715 dose-response in PL$^{pro}$, deubiquitinase, and ISG15 assays*. The $IC_{50}$ (μM) for the (A) PL$^{pro}$,(B) deubiquitinase, and (C) ISG15 assays for DDL-713 and DDL-715 are 54, 51, and 26; and 24, >100, and 39, respectively. The legend in (C) applies to all panels.

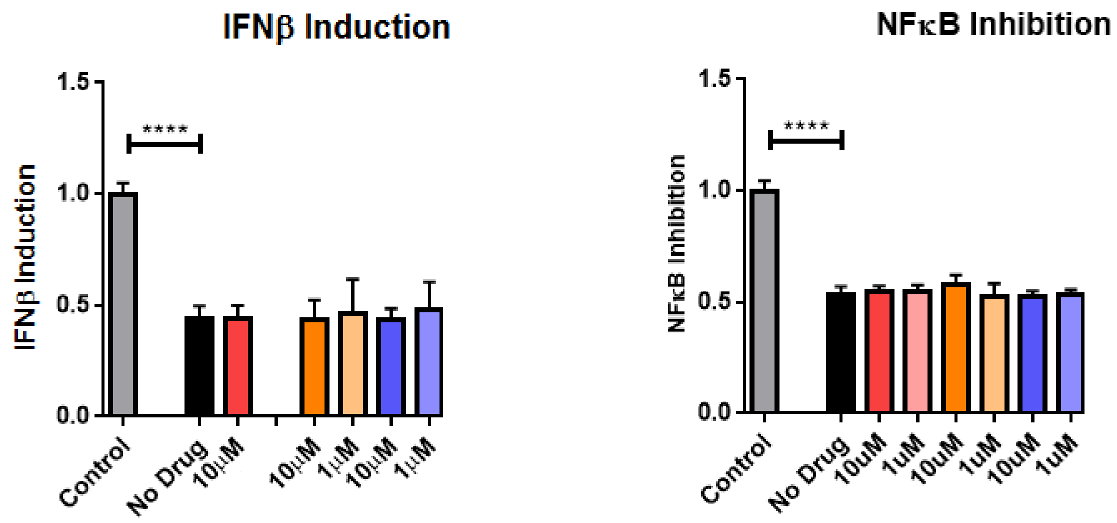


| | Control | No Drug | 10μM | 10μM | 1μM | 10μM | 1μM |
|---|---|---|---|---|---|---|---|
| DMSO (0.5%) | + | + | + | + | + | + | + |
| PLPro | - | + | + | + | + | + | + |
| poly I:C | + | + | + | + | + | + | + |
| 701 | - | - | + | - | - | - | - |
| 715 | - | - | - | + | + | - | - |
| 750 | - | - | - | - | - | + | + |

| | Control | No Drug | 10uM | 1uM | 10uM | 1uM | 10uM | 1uM |
|---|---|---|---|---|---|---|---|---|
| DMSO (0.5%) | + | + | + | + | + | + | + | + |
| PLPro | - | + | + | + | + | + | + | + |
| TNFα | + | + | + | + | + | + | + | + |
| 701 | - | - | + | + | - | - | - | - |
| 715 | - | - | - | - | + | + | - | - |
| 750 | - | - | - | - | - | - | + | + |

**Figure S2**. A) The IFN-β induction assay B) *The NFkB assay*. Compounds were tested at the concentrations listed above (x-axis) and as described in *Methods*.

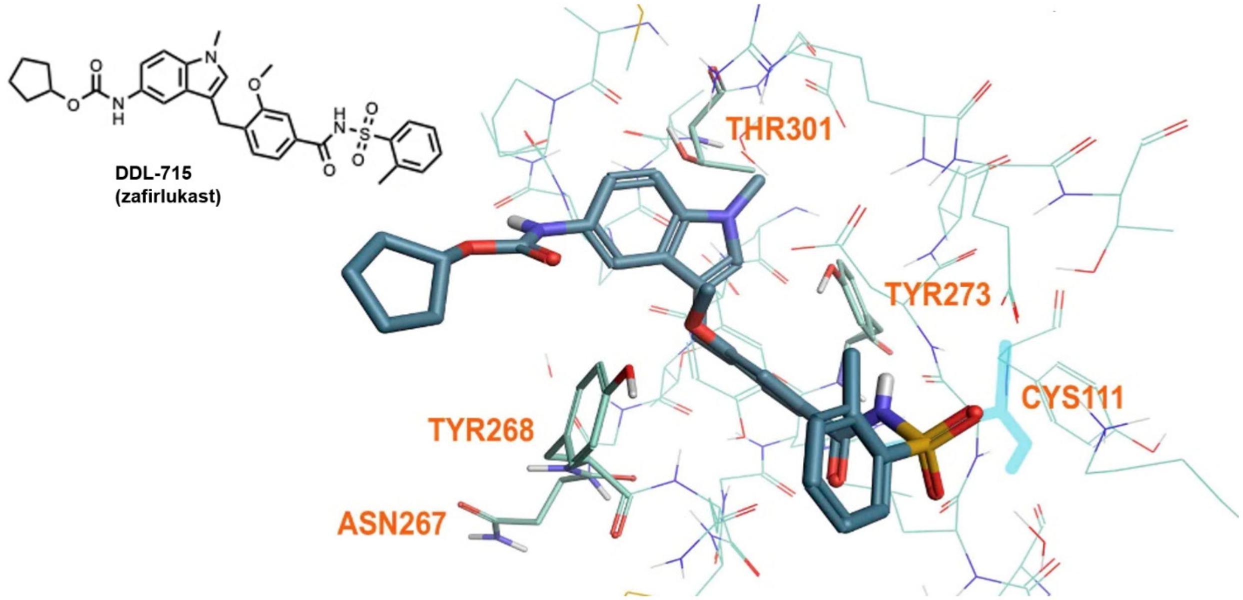


**Figure S3**. Docking of DDL-715 (zafirlukast) using the crystal structure of the PL$^{pro}$ inhibitor GRL-0617 with the SARS-CoV-2 PL$^{pro}$ (PDB ID: 7CMD).

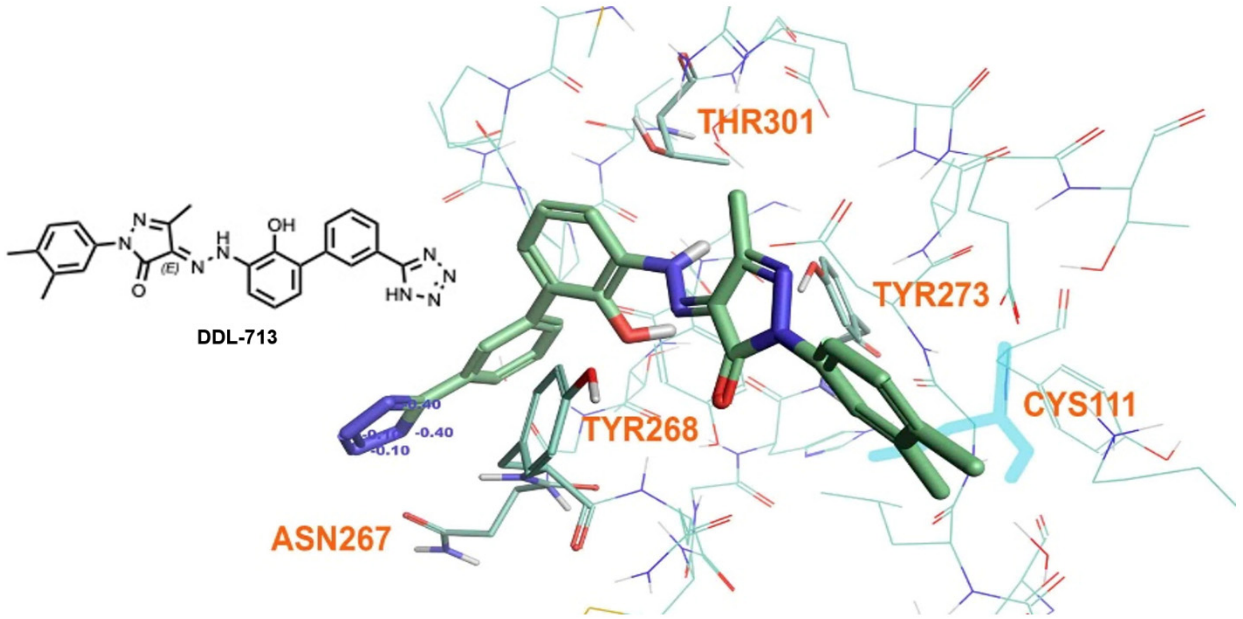


**Figure S4**. Docking of DDL-713 (TPO-agonist-1) using the crystal structure of the $PL^{pro}$ inhibitor GRL-0617 with the SARS-CoV-2 $PL^{pro}$ (PDB ID: 7CMD).

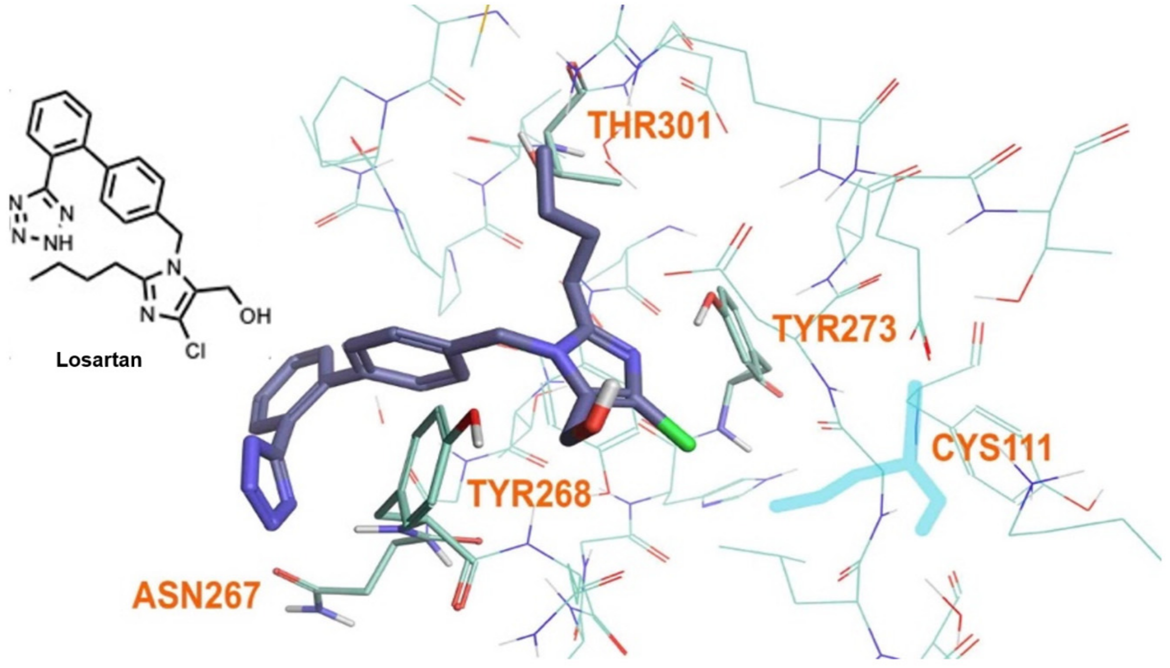


**Figure S5**. Docking of losartan using the crystal structure of the $PL^{pro}$ inhibitor GRL-0617 with the SARS-CoV-2 $PL^{pro}$ (PDB ID: 7CMD).

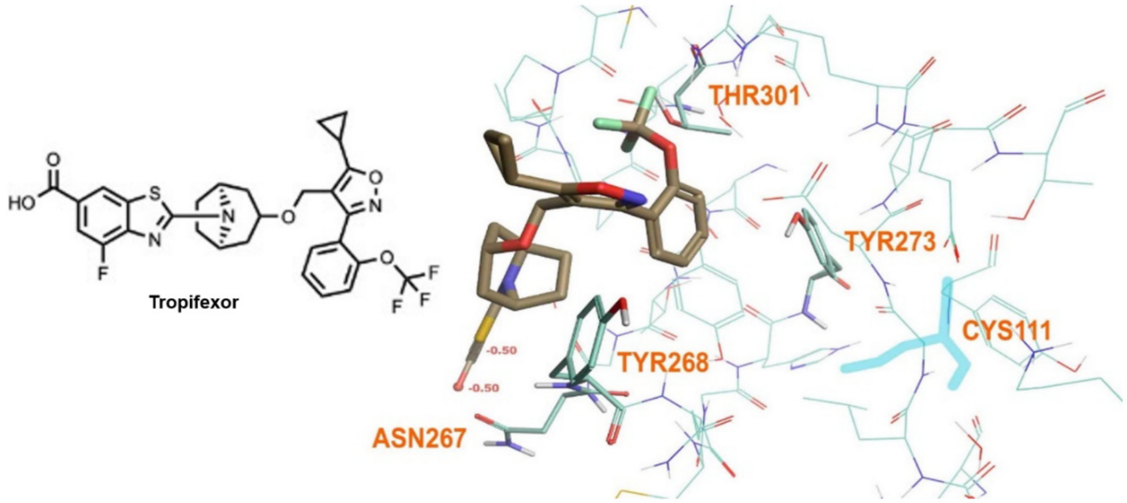


**Figure S6**. Docking of tropifexor using the crystal structure of the PL$^{pro}$ inhibitor GRL-0617 with the SARS-CoV-2 PL$^{pro}$ (PDB ID: 7CMD).

### Supplementary Table

**Table S1.** *The 58-compound custom library screened for PL$^{pro}$ inhibition activity*

| Compound name | CAS | MW | Well# | Pharmacological Class/Activity | Source | Cat # |
|---|---|---|---|---|---|---|
| Abemaciclib | 1231929-97-7 | 540.68 | 1 | CDK inhibitor | 1 Plus Chem | 1P000KLY |
| Aclidinium bromide | 320345-99-1 | 564.6 | 2 | Muscarinic antagonist | Cayman | 20699 |
| Alcaftadine | 147084-10-4 | 307.4 | 3 | H1 receptor antagonist | Cayman | 21290 |
| Avanafil | 330784-47-9 | 483.96 | 4 | PDE5 inhibitor | AKScientific | Z5510 |
| Axitinib | 319460-85-0 | 386.5 | 5 | Tyrosine kinase inhibitor | Cayman | 13813 |
| AZD-5423 | 1034148-04-3 | 430.53 | 6 | Non-steroidal glucocorticoid | MCE | HY-108243 |
| Bazedoxifene | 198481-32-2 | 469.66 | 7 | Selective estrogen receptor modulator (SERM) | MCE | HY-A0031 |
| Berbamine | 478-61-5 | 392.49 | 8 | Calcium channel blocker. | AK Scientific | J98904 |
| Betrixaban | 330942-05-7 | 451.91 | 9 | Factor Xa inhibitor | Sigma | SML2845 |
| Bortezomib | 179324-69-7 | 475.57 | 10 | Proteosome inhibitor | 1 Plus Chem | 1P0027E4 |
| Brigatinib | 1197953-54-0 | 584.09 | 11 | ALK inhibitor | MCE | HY-12857 |
| Budesonide | 51333-22-3 | 470.6 | 12 | Corticosteroid | 1 Plus Chem | 1P00DEL8 |
| Canagliflozin | 842133-18-0 | 444.52 | 13 | sodium-dependent glucose cotransporter 2 (SGLT2) inhibitor | AKScientific | J53622 |
| Capmatinib | 1029712-80-8 | 412.42 | 14 | c-Met kinase inhibitor | MCE | HY-13404 |
| Cisclesonide | 126544-47-6 | 506.59 | 15 | Glucocorticoid | 1 Plus Chem | 1P007307 |
| Cobicistat | 1004316-88-4 | 776.04 | 16 | CYP3A inhibitor | Sigma | ADV465750862 |
| Dasatinib | 302962-49-8 | 488.01 | 17 | Tyrosine kinase inhibitor | Tocris | 6793 |
| Dithiobis(benzothiazole) | 120-78-5 | 332.49 | 18 | Food additive | Sigma | D218154 |
| Diosmin | 520-27-4 | 345.42 | 19 | Glycosylated flavonoid | 1 Plus Chem | 1P00IKXD |
| Disulfiram | 97-77-8 | 296.54 | 20 | Acetaldehyde dehydrogenase inhibitor | Tocris | 3807 |

| | | | | | | |
|---|---|---|---|---|---|---|
| E-64 | 66701-25-5 | 513.5 | 21 | Cysteine protease inhibitor | 1 Plus Chem | 1P0037A9 |
| Ebastine | 90729-43-4 | 610.52 | 22 | H1 receptor antagonist | 1 Plus Chem | 1P003QDK |
| Eltrombopag | 496775-61-2 | 442.5 | 23 | Thrombopoietin receptor agonist | Cayman | 13247 |
| Encorafenib | 1269440-17-6 | 540.01 | 24 | BRAF inhibitor | MCE | HY-15605 |
| Entrectinib | 1108743-60-7 | 560.6 | 25 | Tyrosine kinase inhibitor | Cayman | 19476 |
| Ertuglifozin | 1210344-57-2 | 436.88 | 26 | Sodium-dependent glucose cotransporter 2 (SGLT2) inhibitor | MCE | HY-15461 |
| Esomeprarzole | 119141-88-7 | 357.41 | 27 | Proton pump inhibitor | 1 Plus Chem | 1P0078EN |
| Ezetimibe | 163222-33-1 | 522.57 | 28 | Cholesterol lowering | 1 Plus Chem | 1P001UD4 |
| Fluspirilene | 1841-19-6 | 441.54 | 29 | Antipsychotic | AK Scientific | SYN5406 |
| Fluticasone | 90566-53-3 | 409.43 | 30 | Corticosteroid | MCE | HY-15234 |
| Fursultiamine | 804-30-8 | 398.54 | 31 | Vitamin B1 derrivative | AKScientific | M514 |
| Gluiquidone | 33342-05-1 | 487.45 | 32 | Pottasium channel antagonist | 1 Plus Chem | 1P0035FH |
| GRL0617 | 1093070-16-6 | 304.39 | 33 | PLpro inhibitor | Tocris | 7280 |
| Indoprofen | 31842-01-0 | 281.31 | 34 | Nonsteroidal anti-inflammatory | 1 Plus Chem | 1P003FB4 |
| Ipratropium | 22254-24-6 | 412.37 | 35 | Anticholinergic | Tocris | O692 |
| Ivacaftor | 873054-44-5 | 527.63 | 36 | CFTR activator | 1 Plus Chem | 1P0039Q3 |
| Ivermectin | 70288-86-7 | 608.54 | 37 | Antiparisitic | 1 Plus Chem | 1P00ICI6 |
| Lansoprazole | 103577-45-3 | 369.36 | 38 | Proton-pump inhibitor | Sigma | L8533 |
| Lapatinib | 231277-92-2 | 581.06 | 39 | Tyrosine kinase inhibitor | Tocris | 6811 |
| Loperamide | 53179-11-6 | 477.05 | 40 | Antidiarrheal | AK Scientific | C598 |
| Lumacaftor | 936727-05-8 | 452.41 | 41 | Protein chaperone | 1 Plus Chem | 1P00379M |
| Lusutrombopag | 1110766-97-6 | 516.6 | 42 | Thrombopoietin receptor agonist | 1 Plus Chem | 1P0090BL |
| Montelukast | 158966-92-8 | 575.68 | 43 | leukotriene receptor antagonists (LTRAs) | AvaChem Scientific. | 2041A |
| Omeprazole | 73590-58-6 | 875.09 | 44 | Proton-pump inhibitor | 1 Plus Chem | 1P003TY2 |
| Oxitropium bromide | 30286-75-0 | 412.32 | 45 | Anticholinergic | Sigma | Y0000709 |
| Pantoprazole | 718635-09-7 | 405.35 | 46 | Proton pump inhibitor | Sigma | P0021 |
| Rabeprazole | 117976-90-6 | 381.42 | 47 | Proton-pump inhibitor | Sigma | SML0476 |
| Roflumilast | 162401-32-3 | 403.21 | 48 | PDE4 inhibitor | Tocris | 6641 |
| Rutin | 153-18-4 | 472.42 | 49 | Bioflavonoid | 1 Plus Chem | 1P00HYK6 |
| Salbutamol | 18559-94-9 | 239.31 | 50 | β2 adrenergic receptor agonist | Sigma | S8260 |
| Siponimod | 1230487-00-9 | 442.5 | 51 | Sphingosine-1-phosphate receptor modulator | 1 Plus Chem | 1P009D0W |
| Ticagrelor | 274693-27-5 | 591.55 | 52 | P2Y12 receptor antagonist | 1 Plus Chem | 1P00I5O1 |
| Tiotropium bromide | 136310-93-5 | 472.42 | 53 | Muscarinic antagonist | AKScientific | A626 |
| TPO agonist 1 | 1033040-23-1 | 438 | 54 | Thrombopoietin (TPO) agonist | MCE | HY-100380 |
| Troglitazone | 97322-87-7 | 386.44 | 55 | Antihyperglycemic agent | 1 Plus Chem | 1P00JO1Y |
| Umeclidinium Bromide | 869113-09-7 | 508.5 | 56 | Muscarinic antagonist | AKScientific | 4140AH |
| Voxelotor | 1446321-46-5 | 337.4 | 57 | Hemoglobin polymerization inhibitor | Cayman | 23933 |
| Zafirlukast | 107753-78-6 | 598.66 | 58 | Leukotriene receptor antagonists (LTRAs) | 1 Plus Chem | 1P007EAT |
